# The human oral microbiome is shaped by shared environment rather than genetics: evidence from a large family of closely-related individuals

**DOI:** 10.1101/131086

**Authors:** Liam Shaw, Andre L. R. Ribeiro, Adam P. Levine, Nikolas Pontikos, Francois Balloux, Anthony W. Segal, Adam P. Roberts, Andrew M. Smith

## Abstract

The human microbiome is affected by multiple factors, including the environment and host genetics. In this study, we analyzed the oral microbiome of an extended family of Ashkenazi Jewish individuals living in several cities and investigated associations with both shared household and host genetic similarities. We found that environmental effects dominated over genetic ones. While there was weak evidence of geographic structuring at the level of cities, we observed a large and significant effect of shared household on microbiome composition, supporting the role of immediate shared environment in dictating the presence or absence of taxa. This effect was also seen when including adults who had grown up in the same household but moved out prior to the time of sampling, suggesting that the establishment of the oral microbiome earlier in life may affect its long-term composition. We found weak associations between host genetic relatedness and microbiome dissimilarity when using family pedigrees as proxies for genetic similarity. However this association disappeared when using more accurate measures of kinship based on genome-wide genetic markers, indicating that environment rather than host genetics is the dominant factor affecting the composition of the oral microbiome in closely-related individuals. Our results support the concept that there is a consistent core microbiome conserved across global scales, but that small-scale effects due to shared living environment significantly affect microbial community composition.

**IMPORTANCE:** Previous research shows that relatives have a more similar oral microbiome composition than non-relatives, but it remains difficult to distinguish the effects of relatedness and shared household environment. Furthermore, pedigree measures may not accurately measure host genetic similarity. In this study, we include genetic relatedness based on genome-wide SNPs (rather than pedigree measures) and shared environment in the same analysis. We quantify the relative importance of these factors by studying the oral microbiome in members of a large extended Ashkenazi Jewish family who share a similar diet and lifestyle despite living in different locations. We find that host genetics plays no significant role and that the dominant factor is shared environment at the household level. We also find that this effect appears to persist in individuals who have moved out of the parental household, suggesting that the oral microbiome established earlier in life persists long-term.

## Introduction

The human microbiome is the name given to the collected communities of bacteria that live on and in the human body. The oral microbiome is one of the most diverse (1) of any human-associated microbial community (2). The oral microbiome is a causative factor in conditions such as dental caries (3), periodontal disease (4), and halitosis (5), and has also been implicated as a reservoir for infection at other body sites (2) and in the pathogenesis of non-oral diseases, such as inflammatory bowel disease (6). Strictly speaking there is no single ‘oral microbiome’ as its composition is highly heterogeneous across different sites in the mouth (7, 8), but the term is commonly used to encompass all of these. Site-specific microbiomes can be observed in the periodontal sulcus, dental plaque, tongue, buccal mucosa and saliva (9). The salivary microbiome exhibits long-term stability and can be considered as an important reservoir that contains microorganisms from all distinct ecological niches of the oral cavity. Characterizing and understanding the factors defining salivary microbial composition is thus crucial to understanding the oral microbiome (10, 11).

Some factors that are thought to influence the human microbiome include environment, diet, disease status and host genetics (12). The relative importance of these factors for the oral microbiome is still under debate, with the majority of previous studies focusing on the gut microbiome (7-9), although it seems reasonable to assume some potential interaction between the oral microbiome and microbial communities in other parts of the human body including the intestinal tract (10).

There is evidence that genetically related individuals tend to share more gut microbes than unrelated individuals, whether or not they are living in the same house at the time of sampling (13, 14). However, the level of covariation is similar in monozygotic and dizygotic twins, suggesting that a shared early environment may be a more important factor than genetics (13, 15). The effect of co-habitation with direct and frequent contact is greatest when considering the skin microbiome, with a less-evident effect on the gut and oral microbiomes (11).

There is also evidence that genetic variation is linked to microbiome composition across other body sites, including the mouth (12), with a recent genome-wide association study (GWAS) identifying several human loci associated (p < 5 × 10^−8^) with microbial taxonomies in the gut microbiome (16). However, no study to date has incorporated both genetic relatedness as a continuous variable and shared environment into the same analysis of the oral microbiome.

Despite high diversity between individuals, the oral microbiome appears to have little geographic structure at genus level at the global scale (17). Nevertheless, at smaller geographical scales it appears that the environment plays a role. Song et al. studied 60 household units and found that the bacterial composition of dorsal tongue samples was more similar between cohabiting family members than for individuals from different households, with partners and mother-child pairs having significantly more similar communities (18). However, this did not include information on genetic relatedness in addition to family relationships. It appears that household-level differences in the oral microbiome may also apply to genetically unrelated individuals and non-partners, with a similar pattern observed in analysis of saliva samples from 24 household pairs of genetically unrelated individuals, only half of whom were considered romantic couples at the time of sampling (19).

The establishment of the oral microbiome appears to proceed rapidly in the first few years of life, with a notable increase in diversity from 0-3 years (18), especially after the eruption of teeth (20). The oral microbiome also appears stable within individuals over at least a period of 3 months, with a unique ‘fingerprint’ of oligotypes discernible even within a single bacterial genus (21). These two facts suggest that the establishment of a particular oral microbiome composition early in life could potentially persist into adulthood, particularly if external factors such as diet remain fixed.

A recently described large Ashkenazi Jewish family (22) offers an opportunity to investigate the effect of both environment and genetics in closely-related individuals. The availability of host genetic data for this cohort means that we can calculate similarity between individuals based on SNPs, rather than using measures of relatedness from pedigrees that do not precisely correspond to shared genetic content (23). We hypothesized that using this more accurate measure of host genetic similarity could lead to different conclusions about the proportion of shared microbiome composition attributable to genetics compared with previous studies. Furthermore, due to shared cultural practices we can be reasonably confident that environmental factors such as diet and lifestyle are largely controlled for, compared to other studies where they may be significant confounders (17). For this reason, this cohort represents a unique opportunity to compare the oral microbiome within a large number of individuals living in separate locations but nevertheless sharing a similar diet, lifestyle, and genetic background, and to investigate the long-term effect of shared upbringing on oral microbiome composition.

## Results

### Description of cohort

We found 271 phylotypes in the total dataset, all of which were present when considering just Family A. 49 of these phylotypes were present in >95% of individuals within Family A, with the *Firmicutes* the most abundant phyla (Figure 1a) as observed in previous oral microbiome studies (15, 24). The most abundant genera were *Streptococcus* (30.4%), *Rothia* (18.5%), *Neisseria* (17.1%), and *Prevotella* (17.1%). Composition of samples was similar between the two families (A and B) and the unrelated controls (Figure 1b). These groupings had a small but significant effect in an analysis of variance (*R*^*2*^=0.015, *p*<0.01) but this is typical of comparisons between such large groups that may differ in an unknown number of confounded variables (e.g. diet, genetics, lifestyle). We concluded that Family A was at the very least a representative sample capturing the majority of the variation present in the wider Ashkenazi Jewish population, if not also non-Ashkenazi-Jewish individuals (for comparison with Human Microbiome Project data see Supplementary Figure 4).

**Figure 1.**
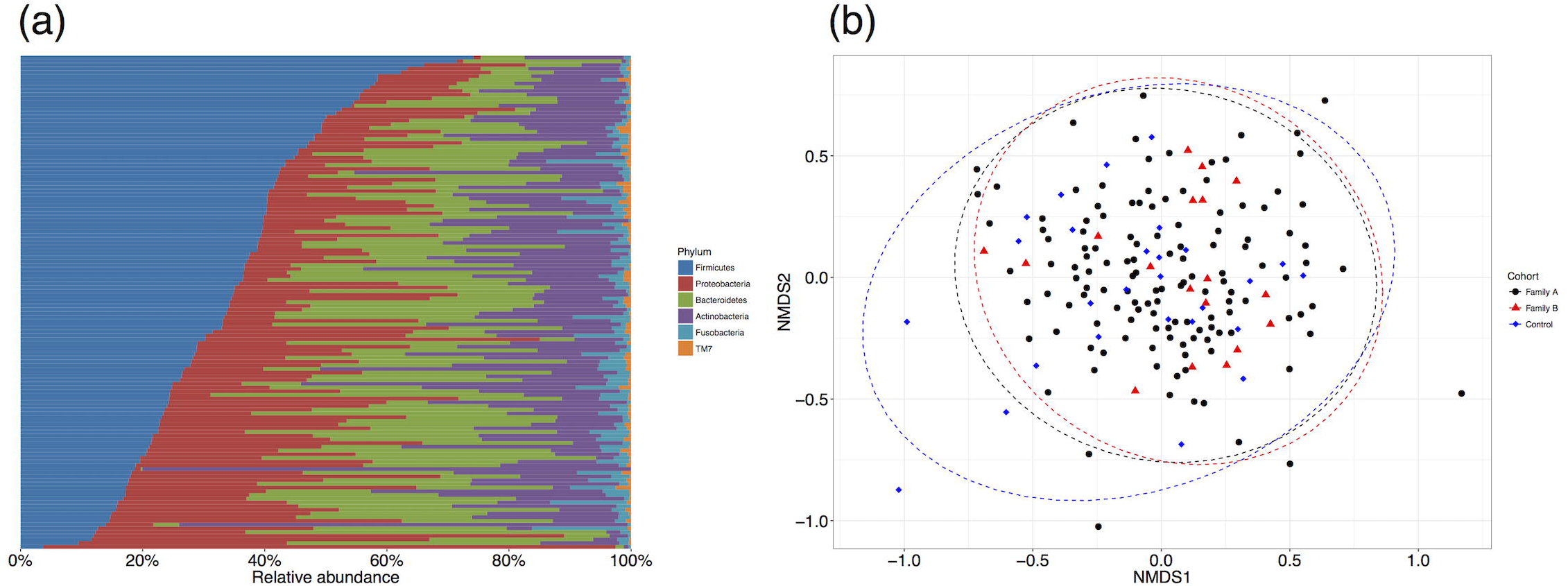
This cohort contains a representative sample of variation in oral microbiome composition. **(a)** Relative abundance of the six bacterial phyla found in saliva samples from Family A, sorted by decreasing *Firmicutes* content. Color scheme adapted from Stahringer et al. (15). Taxonomy was assigned to 271 MED phylotypes using RDP based on the HOMD database. **(b)** Non-metric multidimensional scaling based on Bray-Curtis distances between samples shows high overlap between Family A (black circles), Family B (red triangles), and unrelated Ashkenazi Jewish controls (blue diamonds).

This cohort was originally collected for a study of the genetics of Crohn’s disease (22), and 28 individuals within our sample had a diagnosis of the disease at the time of saliva sample acquisition. We found no significant effect of Crohn’s disease on oral microbiome composition with an exploratory analysis of variance (*R*^2^=0.009, *p*=0.101, *n*=148) accounting for other variables. It was therefore not included as a covariate in further analysis.

### Host genetic similarity is weakly correlated with oral microbiome similarity

We performed an exploratory analysis on individuals in Family A with both genetic and microbiome data available (*n*=111, Supplementary Figure 5), and found that genetic kinship was weakly but significantly associated with oral microbiome dissimilarity computed using Bray-Curtis distances (Supplementary Figure 6; Mantel test *r*=0.065, *p*=0.001). This analysis does not take into account confounding by shared environment, and therefore sets a probable upper bound on the variation that can be attributed to host genetics. Household appeared to be related to oral microbiome composition (Figure 2), but is obviously correlated with variation in host genetics (Supplementary Figure 7) because parents tend to live with their children. This emphasizes the need for a quantitative approach looking at the effect of both household and genetics simultaneously.

**Figure 2.**
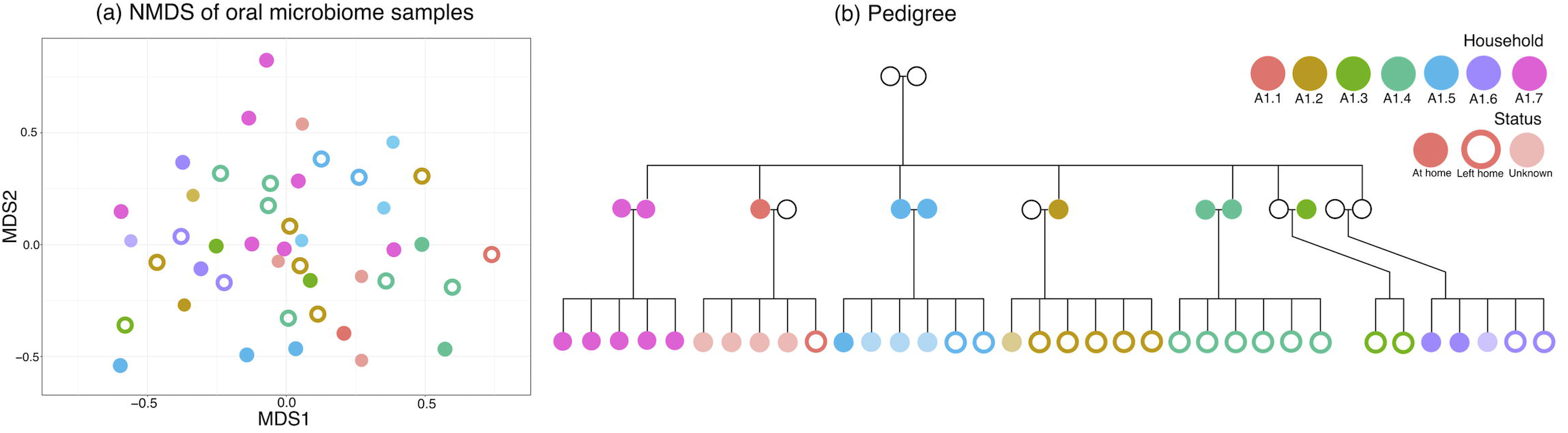
Oral microbiome composition is associated with household. Oral microbiome samples cluster by household (colours), shown by (a) a non-metric multidimensional scaling based on Bray-Curtis distances between samples from individuals in a particular subfamily (*n*=44) within Family A. This figure includes individuals who are currently living together (filled circles), those who had moved out of their childhood home (empty circles), and those for whom data was missing (faint circles). This clustering could be due to shared environment or also due to shared genetics, as is obvious from (b) the pedigree.

### Shared household is the dominant factor affecting oral microbiome composition

We next performed a permutational analysis of variance on the oral microbiome dissimilarities for 28 individuals within Family A, each of whom lived in a household with at least one other individual in the cohort. At the time of sampling, these co-habiting individuals lived across a total of 16 households in four cities (I, II, III, IV). To account for host genetics, we included axes from a metric multidimensional scaling (MDS) of pairwise genetic distances between individuals as explanatory variables (see Methods and Supplementary Figure 8).

There was no significant effect of any of the MDS axes, suggesting that host genetics in closely-related individuals does not significantly affect microbiome composition. We investigated the effect of environment using two levels of geography: city and household (Table 1a). A city-only model showed no significant effect of environment (*R*^2^=0.08, *p*=0.4), whereas a household-only model showed a significant effect (*R*^2^=0.30, *p*=0.001). This was reproduced in a model containing both geographic variables, with permutations stratified by city, where household was still a significant effect (*R*^2^=0.22, *p*=0.001), suggesting that differences at the level of household are more important than at larger geographical scales. We confirmed that city-level effects were small by extending our sample to 82 individuals across the four cities who were not necessarily cohabiting with others (I: 48, II: 13, III: 12, IV: 9), and found that city still had a small effect, although it was significant (*R*^*2*^=0.053, *p*<0.01).

**Table 1.**
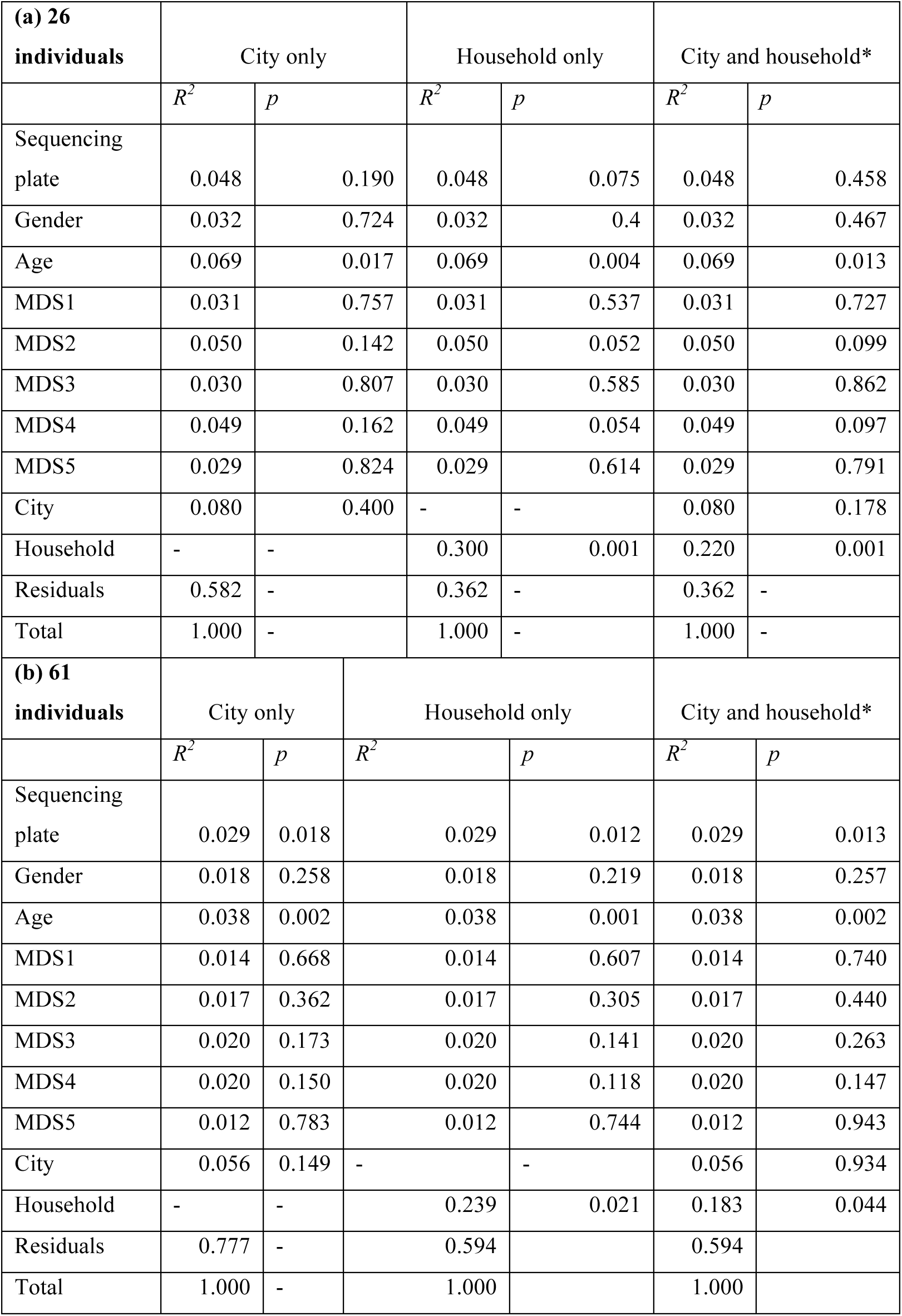
Permutational analysis of variance (adonis) results based on (a) 26 co-habiting individuals who lived in the same household as at least one other individual and (b) 61 individuals who had at least co-habited at some point. *Permutations stratified by city in this analysis.

In this analysis we also found no significant effect of genetics, but age was significant (*R*^*2*^=0.028, *p*=0.0101) (Supplementary Table 1).

### Spouses share taxa at the species level

Restricting the analysis to only married couples within Family A (*n*=16, eight couples), shared household explained even more of the variance (*R*^*2*^=0.591, *p*=0.001). Subtle variations in the relative abundance of phylotypes within the same genus between households were observable, even within the same city location. For example, *Leptotrichia* phylotypes varied consistently between spouse pairs and these patterns were also seen in children living at home (Figure 3). MED phylotype X2772 was present in both spouses in household A1.7, and was also present in the two youngest children within that household (aged 10 or under). Similarly, within household A2.4 the two children aged 10 or under were more similar in *Leptotrichia* phylotypes than an older child. Similar patterns with spouses were also visible in other abundant genera (Supplementary Figure 9).

**Figure 3.**
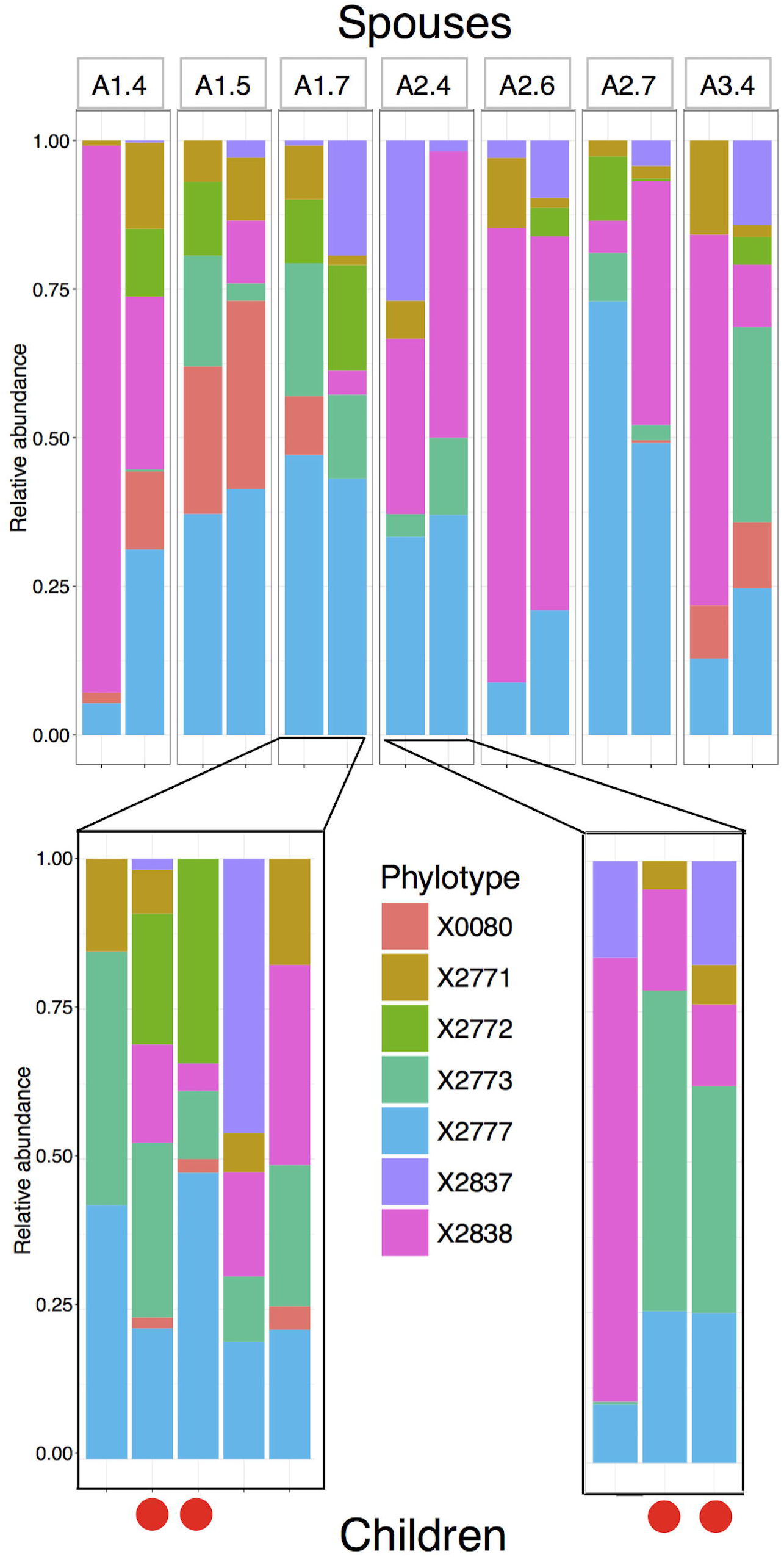
Household-level variation within a genus, shown here with the relative abundance of phylotypes within *Leptotrichia*. The relative abundance of phylotypes within seven pairs of spouses shows clear associations with household. These patterns are to some extent recapitulated in their children. Looking at children still living at home, MED phylotype X2772 is not observed in any individual from household A2.4, but is found in both spouses and two children living in household A1.7. Red dots indicate children aged 10 or under at time of sampling, who appear more similar to each other than other pairs of children. For variation within the top twelve most abundant genera between spouses, see Supplementary Figure 9.

### Household effects persist in individuals who are no longer co-habiting

There were an additional 35 individuals who had grown up in a household with at least one other individual present, but who no longer lived together at time of sampling. To see if the effects of household persisted, we repeated analysis of variance with these individuals included along with the cohabiting-individuals (*n*=61, Table 1b). The effect of household remained significant (*R*^*2*^=0.183, *p*=0.044), and no axes of human genetic variation were significant (*p*>0.05). Age had a significant effect (*R*^*2*^=0.038, *p*<0.01).

Other variables such as age and sequencing plate had smaller effects than household in all our analyses of variance. However, the order of variables can have an effect when performing adonis with an unbalanced design. To check this was not biasing our results, we randomly permuted the order of variables in our model and confirmed that household was always significant (*q*<0.05, Benjamini-Hochberg multiple testing correction) as was age (see Supplementary Material).

### Relying on pedigree kinships produces a genetic signal

To test whether our conclusions required using kinships estimated from genome-wide SNP data for individuals, or whether pedigree information was sufficient, we also repeated our analyses using pedigree kinships (see Methods). Using pedigree kinships resulted in a small but significant amount of variation in microbiome composition being attributable to host genetics via MDS axis 4 (*R*^*2*^=0.016, *p*<0.01, Table 2).

**Table 2.** Comparison of pedigree-based and genome-wide measures of kinship to take host genetics into account in a permutational analysis of variance (adonis) on oral microbiome dissimilarities of *n*=111 individuals. Using pedigree information to produce kinship results in a significant association with human genetics via the fourth MDS axis, which is not present using kinships calculated with LDAK based on genome-wide SNPs.

|  | Pedigree (kinship2) |  | Genome-wide SNPs (LDAK) |  |
| --- | --- | --- | --- | --- |
| | $R^2$ | $p$ | $R^2$ | $p$ |
| Sequencing plate | 0.028 | <0.001 | 0.028 | <0.001 |
| Gender | 0.011 | 0.094 | 0.011 | 0.096 |
| Age | 0.023 | <0.001 | 0.023 | <0.001 |
| MDS1 | 0.010 | 0.174 | 0.011 | 0.119 |
| MDS2 | 0.007 | 0.706 | 0.010 | 0.231 |
| MDS3 | 0.012 | 0.063 | 0.011 | 0.131 |
| MDS4 | 0.016 | 0.009 | 0.011 | 0.111 |
| MDS5 | 0.009 | 0.325 | 0.007 | 0.617 |
| Parental household | 0.215 | <0.001 | 0.217 | <0.001 |
| Residuals | 0.670 | - | 0.671 | - |
| Total | 1 | - | 1 | - |

## Discussion

We have conducted, to our knowledge, the first simultaneous investigation of the role of environment and host genetics in shaping the human saliva microbiome in a cohort of closely-related individuals within a large Ashkenazi Jewish family. We found a weak correlation between host kinship and oral microbiome dissimilarity before taking shared household into account, and an apparent small but significant effect of genetics when using kinships based on the family pedigree as proxies for genetic similarity. However, when using kinship estimates based on genome-wide SNPs between individuals and simultaneously controlling for shared household with a permutational analysis of variance, we find no support for any clear effect of human genetics, suggesting that shared environment has a much larger effect than genetics and is the dominant factor affecting the oral microbiome (*R*^*2*^>0.18). We also observe that shared household explains more variation for spousal pairs than for children, and that younger children living in the same household share subtle variations in phylotype abundance within genera with their parents (Figure 3). Taken together, these observations support the view that human genetics does not play a major role in shaping the oral microbiome, at least not in individuals of the same ethnicity, compared to the environment.

Our results confirm the seemingly paradoxical situation that the oral microbiome is largely consistent across global geographical scales, but can show large variation between households in the same city. Previous studies have also found evidence of small variations in oral microbiome composition comparing samples across a global scale (17). As noted previously, this variation could be influenced by differences in environmental or cultural factors, in which case controlling for these differences would decrease the amount of geographical variation. All individuals in our study follow a traditional Ashkenazi Jewish lifestyle and subsequently are thought to share a similar diet and lifestyle regardless of geographic location (25), which may reduce the variation attributable to city-level differences.

The establishment of the oral microbiome early in life may lead to the persistence of a similar composition over several years. The oral microbiome has been previously observed to be remarkably persistent within individuals over periods of months (21) to a year (26), and we see similar strain-level variation between spouses and their young children (Figure 3). Our results suggest that the oral microbiome composition established early in life via shared upbringing is able to persist for at least several years, because of the persistence of household effects in individuals no longer co-habiting. It has been observed that monozygotic twins do not have significantly more similar gut microbiomes than dizygotic twins (13). Stahringer et al. observed the same effect in the oral microbiome, and also found that twins’ oral microbiomes became less similar as they grew older and ceased cohabiting, concluding that ‘nurture trumps nature’ in the oral microbiome (15). Our findings from a large number of related individuals rather than twins support this view, including the persistence of shared upbringing effects. Shared upbringing appears to be the dominant factor affecting microbiome composition in both the gut and the mouth, rather than genetic similarity. This may have implications for understanding the familial aggregation of diseases such as inflammatory bowel disease, which has been suggested to have an environmental component (27).

The oral microbiome appears far more resilient to perturbation compared to the gut microbiome, with a rapid return to baseline composition after a short course of antibiotics (28). While this could be because of the pharmacokinetics of the antibiotics involved, Zaura et al. speculate that this difference may be due to the oral microbial ecosystem’s higher intrinsic resilience to stress, as the mouth is subject to more frequent perturbation (29). Our work supporting the dominant role of the environment in affecting oral microbiome composition suggests that another important factor in long-term persistence may be the regular reseeding of the ecosystem with bacteria from the external environment.

The fact that we reached our conclusion about the lack of effect of genetics only after including kinship based on genome-wide SNP markers casts doubt on the reliability of pedigrees for calculating relatedness. There are several possible reasons for a discrepancy between kinship estimates from pedigrees and allele sharing (23). One possibility is errors in the pedigree, most likely due to extra-pair paternities, although this explanation can be ruled out in this dataset. More importantly, inherent stochasticity in the Mendelian process of inheritance means that although parents always pass on 50% of their genes to their offspring, SNPs are inherited together in blocks (i.e. haplotypes), meaning that the relatedness between two offspring in a family can be substantially different from 50%. Finally, and most importantly for this closely-related population, shallow pedigrees cannot fully capture complex inbreeding patterns. Thus, while pedigrees are a good model for host relatedness in microbiome studies of large randomly mating populations, they should be used with caution in closely-related large families like this one.

### Limitations

Because all individuals in our main cohort were members of the same extended Ashkenazi Jewish family, the genetic variation in our dataset is therefore much lower than between individuals from a wider population. It is conceivable that host genetics between more distantly-related individuals may play a significant role in affecting oral microbiome composition. Furthermore, our results only looked at overall genetic similarity, assessed using community comparison metrics based on taxa abundances. They therefore do not preclude the existence of fine-scale links between particular microbial taxa and individual genetic loci, particularly in immune-sensing genes such as those identified in the gut microbiome by Bonder et al. using a much larger cohort (30), although our study was not designed or powered to detect such associations.

Additionally, we lack detailed information on diet and lifestyle factors of individuals in this study. However, the shared cultural practices within this Ashkenazi Jewish family mean that it is not unreasonable to assume they share similar lifestyles and diet despite living in different locations around the world (25).

The apparent persistence of shared upbringing could be confounded by the fact that individuals may continue living near to the household where they grew up. If this were the case, then our observation could instead be due to the persistence of a shared environment beyond the household at a level intermediate between household and city, rather than the persistence of a stable oral microbiome following environmental change. Finally, our samples represent only a single cross-sectional snapshot in time. More long-term longitudinal studies like the work of Stahringer et al. on twins (15) are necessary to investigate the persistence of the oral microbiome after its establishment early in life in a variety of relatedness settings.

## Conclusion

In summary, our results incorporating a measure of genetic relatedness using SNPs demonstrate that the overall composition of the human oral microbiome in a large Ashkenazi Jewish family is largely influenced by shared environment rather than host genetics. An apparent significant effect of host genetics using pedigree-based estimates disappears when using genetic markers instead, which recommends caution in future microbiome research using pedigree relatedness as a proxy for host genetic similarity. Geographic structuring occurs to a greater extent at household level within cities than between cities on different continents. Living in the same household is associated with a more similar oral microbiome, and this effect persists after individuals have left the household. This is consistent with the long-term persistence of the oral microbiome composition established earlier in life due to shared upbringing.

## Materials and Methods

### Ethics

Ethical and research governance approval was provided by the National Research Ethics Service London Surrey Borders Committee and the UCL Research Ethics Committee. Written informed consent was provided by all participants.

### Cohort

Our cohort contained data from 133 individuals within the same extended family (Family A) living in four disparate cities (I, II, III, IV) across three continents (see (22) for more information). We also had samples available from 18 individuals from a separate smaller family (Family B), and 27 unrelated Ashkenazi Jewish controls. All individuals studied were of genetically confirmed Ashkenazi Jewish ancestry (22, 25). When not directly available, shared household was inferred according to age; individuals within this community marry and subsequently leave the family home at a median age of 21 (95% CI: 19-26) (25). Therefore, we assumed that individuals aged 18 or younger at the time of sampling were living with their parents and individuals aged 25 or older were not.

For analysis of the effects of household, we included only households with two or more individuals so as to remove the possibility that we were only measuring inter-individual differences, which can be large in the oral microbiome (17, 21). 26 individuals were living with at least one other individual at the time of sampling in a total of nine households. An additional 35 individuals who had grown up in a shared household with at least one other individual in the cohort, but who were no longer living together were subsequently included in the analysis.

### Sampling

Saliva samples were collected in sterile tubes containing saliva preservative buffer as per the method of Quinque et al. (31). For full protocol see the Supplementary Material. 500ml of saliva/preservative buffer were used with PurElute™ Bacterial Genomic Kit (Edge Biosystems, Gaithersburg, MD) for DNA extraction. After bacterial DNA extraction, three spikes were added to all samples in a final concentration of 4pg/ml, 0.4pg/ml and 0.08pg/ml, respectively.

### PCR amplification, purification and sequencing

The Mastermix 16S Basic containing MolTaq 16S DNA polymerase (Molzym GmbH & Co.KG, Bremen, Germany) was used to generate PCR amplicons. PCR amplicons were purified in two rounds using the Agencourt® AMPure® system (Beckman Coulter, Beverly, Massachusetts) in an automated liquid handler Hamilton StarLet (Hamilton Company, Boston, Massachusetts). DNA quantitation and quality control was performed using the Agilent 2100 Bioanalyzer system (Agilent Technologies, Inc., Santa Clara, CA). We used 785F and 1175R 16S rRNA primers (Supplementary Data Table 1) that amplified the V5-V7 region of the 16S rRNA gene on the Illumina MiSeq System (Illumina, San Diego, CA).

### Quality control

To assess technical variation across runs, we spiked samples during library preparation with a fixed amount of synthetic DNA (see Supplementary Material). Three unique spike sequences (length 350) were designed which could be easily identifiable for quality control purposes. We found, as expected, that the number of spike sequences and the number of putative 16S sequences (length between 350 and 380 bases) were negatively correlated with each other due to the limited total sequencing depth of the Illumina Miseq (Supplementary Figure 1). The variation in reads corresponding to this spike across samples was independent of run. We also resequenced a subset of samples without spikes to verify whether spikes affected our analyses and observed the same qualitative differences (Supplementary Figure 2), implying that the addition of spikes did not have a negative impact on downstream analysis. Paired-end reads were merged with fastq-mergepairs in VSEARCH v1.11.1 (32), discarding reads with an expected error >1. As the expected length of the V5-V7 region was 369 bases, we discarded sequences with <350 or >380 bases.

### Clustering and taxonomic classification

Sequences were clustered with Minimum Entropy Decomposition (MED) (33). MED requires that the variation in read depth across samples does not differ by several orders of magnitude, so we discarded samples with fewer than 5,000 reads and subsampled to a maximum number of 20,000 sequences, resulting in 6,353,210 sequences. We ran MED v2.1 with default parameters (see Supplementary Methods), identifying 271 phylotypes in the dataset (Supplementary Table 2). MED offers higher resolution compared to Operational Taxonomic Unit (OTU) picking methods, and has previously been shown to differentiate the composition of the oral microbiome of individuals over time even within the same genus (21). We verified that using MED phylotypes gave very similar compositional dissimilarities compared to using OTUs (Supplementary Figure 3) but allowed slightly increased statistical power in analysis of variance (see Supplementary Material), consistent with the literature (33). MED phylotypes had taxonomy assigned using RDP (34) against the Human Oral Microbiome Database (HOMD) (35). Comparison to Human Microbiome Project (HMP) oral samples also indicated that Ashkenazi Jewish individuals do not have a significantly different oral microbiome from other populations, with Ashkenazi Jewish saliva samples clustering with non-plaque samples from individuals in the HMP (Supplementary Figure 4). However, the use of different primers makes it difficult to reach a robust conclusion on this point.

### Inclusion of host genetics

We investigated the effect of relatedness between individuals on oral microbiome composition using both genetic kinships (based on SNPs) and pedigree kinships (based on the pedigree). We calculated pedigree kinships with kinship2 (36) and genetic kinships with LDAK v5.94 (37) using genome-wide SNP data from either the Illumina HumanCytoSNPv12 (Illumina, USA) or the Illumina HumanCoreExome-24, as described previously (22). These genetic kinships *K*_g_ are normalized to have a mean of zero, and correspond to genetic similarity between individuals. *K*_g_correlates with the pedigree kinship *K*_g_ but there can be substantial spread around the expected values due to the random nature of genetic inheritance (Supplementary Figure 4a), making *K*_g_ a more accurate measure of true genetic similarity between individuals (23).We converted these kinships to dissimilarities and then Euclidean distances (Supplementary Information) which were used in a multidimensional scaling (MDS) ordination (Supplementary Figure 8).

Following Blekhman et al. (12) we used MDS with five axes as covariates in a permutational analysis of variance of oral microbiome dissimilarities.

### Statistical analysis

We calculated Bray-Curtis dissimilarities between samples based on relative abundances of phylotypes, excluding samples with fewer than 1000 reads. Variance explained in Bray-Curtis dissimilarities was calculated using the adonis function from the vegan v2.4.1 package in R, which performs a permutational analysis of variance of distance matrices (38). We used *n*=9999 permutations, with permutations stratified by geographical sample location where appropriate.

## Funding information

The project was supported through the following charities and research councils: EFFORT (Eastman Foundation For Oral Research and Training), the Charles Wolfson Charitable Trust, and the Medical Research Council (MR/L000261/1). APL was supported by the Irwin Joffe Memorial Fellowship. LS was supported by the Engineering and Physical Sciences Research Council (EP/F500351/1). ALRR was supported by CAPES foundation, Ministry of Education of Brazil (0698130). The funders had no role in study design, data collection and interpretation, or the decision to submit the work for publication.

## Acknowledgments

The authors acknowledge all individuals who kindly participated in this study.

